# Ciliary marginal zone of the developing human retina maintains retinal progenitor cells until late gestational stages

**DOI:** 10.1101/2024.09.03.611053

**Authors:** Kiara C. Eldred, Sierra J. Edgerton, Isabel Ortuño-Lizarán, Juliette Wohlschlegel, Stephanie M. Sherman, Sidnee Petter, Gracious Wyatt-Draher, Dawn Hoffer, Ian Glass, Anna La Torre, Thomas A. Reh

## Abstract

Non-mammalian vertebrates maintain a proliferative stem cell population at the far periphery of their retina called the ciliary marginal zone (CMZ), which gives rise to all retinal cell types and contributes to retinal regeneration upon injury. Humans do not maintain a proliferative CMZ into adulthood; however, it is not known how long in development this region continues to generate new neurons. Here, we identify a population of cells in the far peripheral retina of the fetal human that continues to proliferate long after the rest of the retina is quiescent. Single cell RNA-sequencing and EdU tracing at late time points in development reveal that this region has features of the non-mammalian CMZ, including the capacity to produce both early and late born cell types at late developmental stages, and a longer cell cycle than more centrally located retinal progenitor cells (RPCs). Moreover, while more central RPCs exit the cell cycle with the addition of a TGFβ-inhibitor, we show that early RPCs within the CMZ do not. These findings define the late stages of neurogenesis in human retinal development, and present a unique model system to study the fetal CMZ in humans.

## Introduction

The development of the retina is highly coordinated temporally and spatially, producing a stereotypic structure with a diverse complement of neuronal subtypes^1^. Most cell types within the retina arise from a common progenitor pool^2^. These retinal progenitor cells (RPCs) produce specific subtypes of retinal cells in temporal windows, with early-born cell types produced first, and late-born cell types produced later in development^3,4^. Careful observations of RPCs over time have allowed for the identification of genes that are enriched early RPCs, and a different subset that are more highly expressed in late RPCs^5,6^. Studies focused on human RPCs have identified early RPCs from 63 to 98 days gestation, and seen that late RPCs predominate from 90-190 days gestation^6^. However recent studies have identified late born Müller glia in the central retina as early as day 59 of fetal development, suggesting that a small population of late RPCs must be present much earlier^7^.

Although much of retinal histogenesis occurs during embryogenesis, in many non-mammalian organisms such as the zebrafish, frog, and chick, there is a region of retinal stem cells in the far periphery of the retina that continues to proliferate and generate new neurons post-embryonically^8,9^. This zone of continued neurogenesis is at the border between the neural retina and the ciliary epithelium, and has been termed the Ciliary Marginal Zone (CMZ)^9,10^. The CMZ is a particularly interesting region, as the RPCs maintained there are competent to make all retinal cell types. Proliferation in this region is not believed to be present in adult mammals, and studies in adult mice and humans have failed to find conclusive *in vivo* evidence for neurogenesis at an equivalent zone^10,11^. Recent reports in mice suggest that in the developing retina, a CMZ-like zone with proliferating RPCs, exists transiently at the end of retinal neurogenesis^12^. Moreover, recent studies have reported a CMZ-like zone in the fetal human retina^13^. These studies suggest that the proliferating cells in the human fetal CMZ have markers of early progenitors; however, the RPC populations of the far peripheral human retina have only been well-documented up to 91 days of gestation^13^ due to technical limitations of spatial single cell assays, difficulty of isolating this region, low tissue availability, and the small amount of CMZ tissue present for single cell processing. Therefore, there is not much known about the similarities of the fetal human CMZ and the analogous zone in other species, particularly at later time points in development.

Here, we performed histological analyses of late stages of the CMZ in human fetal development from over 40 human fetal retinas, 4 macaque retinas, and implemented a new culturing method to identify and track the latest proliferating cells of the human retina. We confirmed these RPCs are located in the analogous region to the CMZ of other species. We further characterized these RPCs through single cell RNA-sequencing at 185 days equivalent gestation, identifying populations with early and late progenitor gene expression. We went on to show that both early and late born cell types (ganglion cells versus rods and Müller glia, respectively) are generated at this zone as late as 170-220 days equivalent gestation. We also observed that RPCs of the CMZ are unique in cell cycle length, with a longer cell cycle than those in other regions of the retina, consistent with the CMZ of other species^14,15^. Through EdU labeling at late timepoints in development, we revealed the presence of RPCs until 350 days equivalent development, which is comparable to 2.5 months postnatal development. These *in vitro* results were confirmed by staining in fetal tissues up to day 238 of human retinal development, and also in non-human primates at 145 days gestation, which is equivalent to 263 days human gestation. Furthermore, we showed that late RPCs in human fetal retinospheres prematurely exit the cell cycle with the addition of TGF-β inhibitor, whereas early RPCs of the CMZ are unaffected by inhibition of this signaling pathway. These findings define the end of neurogenesis in human retinal development and provide a new *in vitro* system to further study the biology of the human CMZ.

## Results

### The far peripheral human fetal retina maintains a compact zone of retinal progenitors

The CMZ of non-mammalian vertebrates lies between the laminated retina and the ciliary epithelium at the retinal margin. Therefore, to identify the analogous zone in humans at fetal stages, we examined the peripheral retina at Day 120 of fetal development. We found that the far periphery of the fetal retina, adjacent to the ciliary body, is distinct from the adjacent peripheral retinal tissue in that it lacks lamination of differentiated retinal cells (Fig. 1A-C). To determine if this region contains RPCs, we stained for two transcription factors, VSX2 and PAX6, that are co-expressed in RPCs of the fetal retina as well as in the CMZ in non-mammalian species^8^. At 120 days of fetal development, there are no VSX2/PAX6 co-labeled RPCs in the laminated retina, including the laminated peripheral retina (Fig. 1B). Instead VSX2 labels bipolar cells and PAX6 labels amacrine cells and ganglion cells (Fig. 1B). However, in the far peripheral retina, there are identifiable RPCs that co-express VSX2 and PAX6 (Fig. 1C, empty arrowhead). In addition, a distinct zone lying between the ciliary body and the retina contains cells that co-express both VSX2 and PAX6 in higher amounts than the RPCs in the laminated retina (Fig. 1C, filled arrowhead). The expression of VSX2 and PAX6 in this zone is similar to the expression that occurs in the CMZ of birds^8^.

**Figure 1:**
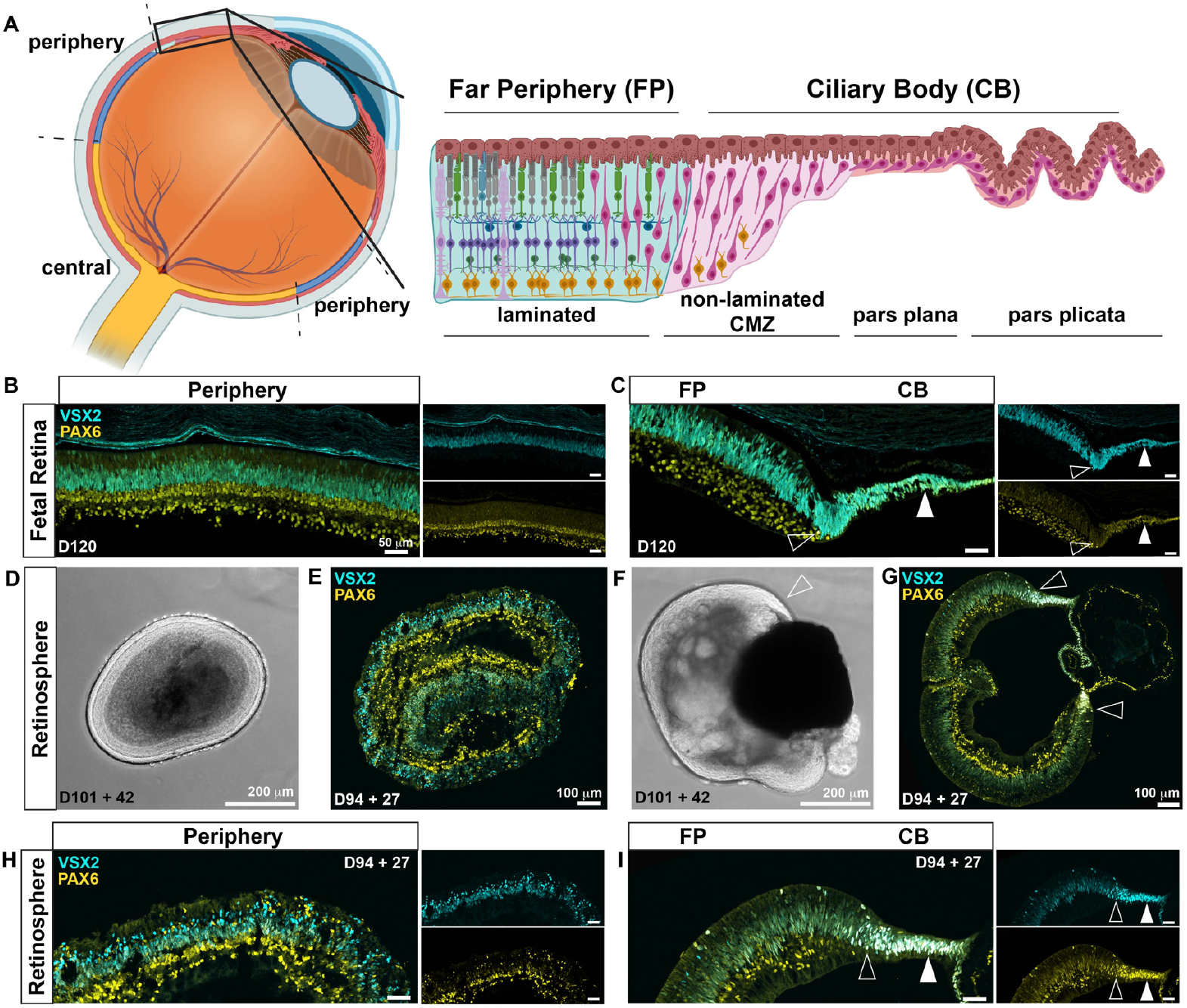
Retinospheres maintain laminated retina and CMZ. **(A)** Schematic of the human retina. Central (yellow) and peripheral (blue) regions. At the far peripheral edge, enlarged region shows far peripheral retina (cyan), and directly adjacent the non-laminated ciliary marginal zone (CMZ, pink). Brown cells lining the top of the enlarged image are retinal pigmented epithelium. **(B-C)** Fetal retina, isolated and fixed at day 120 of gestational development. Immunostaining for VSX2 (cyan) and PAX6 (yellow). Colocalization of these proteins (cyan + yellow = white) indicates progenitor cells. Scale bars are 50 μm. **(B)** Peripheral region. No obvious colocalization of VSX2 and PAX6 is present. **(C)** Far Peripheral (FP) region of the fetal retina. Colocalization of VSX2 and PAX6 is observed in the CMZ (open arrowhead) and at the thinning ciliary body (CB, filled arrowhead). **(D)** Peripheral retinosphere isolated at 101 days gestation and cultured for 42 days, 143 days equivalent gestational development, imaged with brightfield. **(E)** Full section of peripheral retinosphere isolated at 94 days gestation and cultured for 27 days, 121 days equivalent gestational development. Immunostaining for VSX2 (cyan) and PAX6 (yellow). **(F)** Retinosphere containing FP and CB, isolated at 101 days gestation and cultured for 42 days, 143 days equivalent gestational development, imaged with brightfield. Open arrowhead indicates CMZ region. **(G)** Full section of FP and CB containing retinosphere, isolated at 94 days gestation and cultured for 27 days, 121 days equivalent gestational development. Immunostaining for VSX2 (cyan) and PAX6 (yellow). Open arrowhead indicates CMZ regions. **(H)** Peripheral retinosphere from Fig. 1E, no obvious colocalization of VSX2 and PAX6 is present. **(I)** Retinosphere containing FP and CB from Fig. 1G, colocalization of VSX2 and PAX6 is observed in the CMZ (open arrow) and at the thinning ciliary body (CB, filled arrow).

To determine how long the progenitor cells persist in the peripheral retina and at the retinal margin, it is necessary to examine older stages of development. Because availability of fetal tissues over 140 days of gestational development is limited, we employed a method to culture fetal retinal tissues for extended periods of time. Previously our lab has shown that the developmental timing of dissected regions of fetal retina cultured as free-floating spheres, termed retinospheres, closely follows fetal development when maintained in culture^5^. Retinospheres were generated from the retina of a 94 day gestation fetus, and derived from either the peripheral retina, or the far peripheral (FP) retina which contains the CMZ region. Retinospheres were cultured for 27 days, to an equivalent gestational age of 121 days. Retinospheres generated from either the peripheral laminated retina (Fig. 1D,E), or the FP and CMZ containing region (Fig. 1F,G) maintain laminar arrangement of retinal cells after isolation and those from the FP show a distinct CMZ region after 27 days *in vitro*, to an equivalent of 121 days of gestation.

The laminated peripheral retinospheres isolated from day 94 and cultured to the gestational equivalent timepoint of 121, a similar timepoint as the fetal tissue described above that was isolated directly (Fig. 1B), do not contain VSX2/PAX6 co-labeled progenitor cells (Fig. 1H). By contrast the FP retina containing the CMZ from the same experiment contain RPCs that coexpress both VSX2 and PAX6 (Fig. 1I). When we examine even later stages, either with or without culture, we see similar distributions of PAX6 and VSX2 in human fetal samples obtained from day 238 fetal retina (Fig. S1A), and retinospheres isolated at day 103 and cultured for 246 days to a total of 249 days gestational equivalent (Fig. S1B-C). These results show that there is an identifiable CMZ up to day 238 days of human fetal development and confirm that retinospheres are a reliable proxy for fetal retina with regards to RPC markers.

### Cells in the CMZ remain mitotically active after the rest of the retina is no longer proliferating

In order to determine how long the CMZ is mitotically active, we generated retinospheres from both laminated peripheral retina and far peripheral retina (regions displayed in Fig. 1A), then added 5-Ethynyl-2′-deoxyuridine (EdU) to mark newly synthesized DNA in dividing cells during specific time windows. When EdU is added at early developmental timepoints for a period of 7 days to retinospheres from day 101 of gestational development and maintained in the culture media for 7 days until fixing at day 108 (Fig. 2A), we see that EdU marks the PAX6/VSX2+ cells at the periphery of the retina, in both the laminated retina and in the non-laminated CMZ region adjacent to the ciliary body (Fig. 2B). When retinospheres isolated from the same retina are maintained in culture to 146 days equivalent gestation and treated with EdU for the last ten days of development (Fig. 2A), we see that the EdU labeled cells are still maintained at the small region of cells adjacent to the ciliary body, and some co-express OTX2, indicating these proliferating cells have differentiated into photoreceptor or bipolar cells (Fig. 2C). By contrast, retinospheres derived from laminated peripheral retina at 101 days and maintained *in vitro* for the same timeframe of 146 day equivalent gestation, have little to no EdU labeled cells (Fig. 2D). All retinal tissue of the retinospheres with a CMZ contain proliferating cells at day 132, as demonstrated by the global incorporation of EdU throughout a 186 day retinosphere when EdU is added continuously from 132 to 186 of development (Fig. 2E and F). In contrast, when EdU is added late for only two days, from day 184 to 186 of development, proliferation is confined to a much smaller region at the edge of the retinosphere in the CMZ (Fig. 2E, G-G’). Quantification of EdU incorporation into retinospheres over time showed that central and laminated peripheral retinospheres have very little EdU at the late time point of 185 days equivalent gestation, while the far peripheral retinosphere regions continue to incorporate EdU at this time (CMZ= 34 EdU+ cells/mm; FP= 9, p = 0.0269) (Fig. S2A-B). This phenomenon of proliferation concentrated in the CMZ region of the retinosphere is observed in 14 separate fetal samples cultured for varying lengths of time (Fig. S2A).

**Figure 2:**
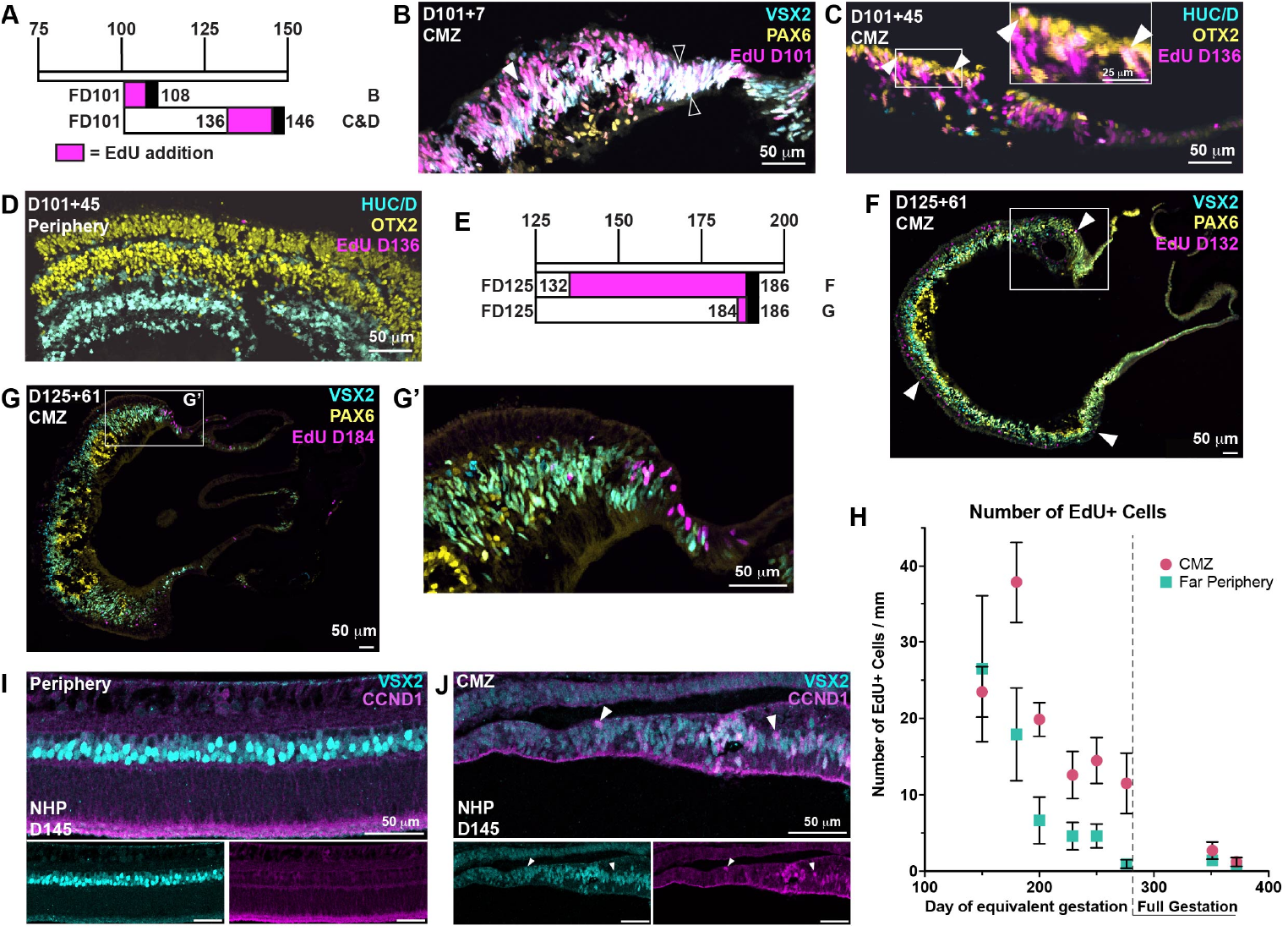
CMZ remains mitotically active until 2.5 months postnatal development. **(A)** Time course of retinosphere isolation, culture, and EdU addition. Magenta signifies EdU addition, black bar signifies final day of fixation. Letters “B”, “C&D” at right correspond to subsequent figures. **(B-D)** Retinosphere containing the late proliferative zone (CMZ), isolated at 101 days gestation and cultured for 7 days, to 108 days equivalent gestational development. **(B)** Immunostained for VSX2 (cyan), PAX6 (yellow), EdU (magenta). Arrows indicate co-expression of all three markers (white), indicating proliferating progenitor cells. Filled arrowhead shows a co-expressing cell in the far periphery, open arrowheads indicate co-expressing cells in the CMZ. **(C)** EdU was added on day 136 until fixing. Immunostained for HUC/D (cyan), OTX2 (yellow), EdU (magenta). Arrows indicate co-expression of OTX and EdU, indicating that these cells have differentiated into a retinal neuron, either a photoreceptor or a bipolar cell. **(D)** EdU was added from isolation on day 136 until fixing. HUC/D (cyan), OTX2 (yellow), EdU (magenta). No appreciable EdU is observed. **(E)** Time course of retinosphere isolation, culture, and EdU addition. Magenta signifies EdU addition. Letters “F” and “G” at right correspond to subsequent figures. **(F-G)** Retinosphere containing the ciliary marginal zone (CMZ), denoted by a box, isolated at 125 days gestation and cultured for 61 days, to 186 days equivalent gestational development. EdU was added on day 132 until fixing. Immunostained for VSX2 (cyan), PAX6 (yellow), EdU (magenta). EdU incorporation can be observed throughout the retinosphere, indicated by filled arrows. **(G)** EdU was added on day 184 until fixing. Immunostained for VSX2 (cyan), PAX6 (yellow), EdU (magenta). EdU incorporation can be observed primarily at the very edge of the retinal tissue in the CMZ. **(G’)** Enlarged image of inset region displayed in G. **(H)** EdU incorporation over time. Number of EdU+ cells were counted in the CMZ region, marked by co-expression of PAX6 and VSX2, and compared to the number of EdU cells in the rest of the retinosphere containing the far periphery. Counts are normalized to mm of retinal length. Error bars displayed are SEM. 14 individual samples quantified for these data are listed in Supplemental Figure S1D, minimum of 2 retinospheres with an average of 4 retinospheres per timepoint. **(I-J)** Non-human primate (NHP) samples from gestational day 145, which is equivalent to 263 days human gestation, immunostained for VSX2 (cyan) and CCND1 (magenta). Peripheral retina. **(J)** CMZ region.

To better characterize the contribution of the human fetal CMZ to retinal growth, we measured the size of retinospheres and the addition of EdU at different time points over the duration of culturing. When CMZ containing retinospheres were measured over time, they continued to increase in size from the day of isolation, either day 101 or day 122, until day 270 equivalent gestation (Fig. S2C-D). This is contrasted by the consistent size of retinospheres isolated from peripheral or central retina regions, which remain a constant size after isolation (For both ages Wallis Multiple Comparison tests; CMZ vs. Periphery p <0.0001, CMZ vs Center p < 0.0001, Periphery vs Center = ns) (Fig. S2C-D). These results indicate that the CMZ cells continue to generate new retinal tissue for up to 270 days of equivalent gestation.

To determine when the CMZ stops proliferating in the fetal human retina, EdU was added for the last three days before fixation, with samples ranging from 170 days equivalent gestation through 370 days, which is the equivalent of 3 months postnatal development (Fig. S2A). We observed EdU incorporation in the CMZ region until 350 days equivalent gestation, which is similar to 2.5 months postnatal development (Fig. 2H). Little to no EdU incorporation was observed in either the laminated peripheral retina or the CMZ past 350 days equivalent development, suggesting that RPCs stop proliferating and have differentiated by this timepoint (Fig. 2H). These results show that the CMZ in the far peripheral retina contains cells that remain mitotically active up to 350 days.

To compare our extensive *in vitro* observations to *in vivo* tissues, we conducted studies in non-human primate (NHP), Rhesus monkey (*Macaca mulatta*) retinal tissues collected at late gestational time points. We use elevated levels of CCND1, which is expressed in cycling cells^16,17^, as an indicator of potential proliferation. At the latest time point available, day 145, Rhesus monkey gestational development is approximately equivalent to day 260 of gestational development in humans^18^. Here we observe lower levels of CCND1 staining in peripheral retinal regions (Fig. 2I); however, a population of CCND1 positive cells is maintained in the CMZ region (Fig. J). The latest evidence we have of proliferation in primary tissues from human fetal stages is from a day 238 fetal sample where Ki67 staining, a marker of cell proliferation, is observed only in the CMZ region and not in the peripheral retina (Fig. S2E-G). These primary tissue studies correlate well with the *in vitro* observations of the CMZ region in retinospheres, and support our conclusion that the CMZ region may persist until birth in the human retina.

### Identification of early and late retinal progenitor cells in the human fetal CMZ

As noted above, retinospheres cultured to late stages of equivalent development have significantly more proliferation when they contain the CMZ compared to those derived from peripheral retina without the CMZ (Fig. 2G, H). To better understand the characteristics of these replicating cells, we compared the cell type composition of peripheral retinal to the composition of the CMZ using single cell RNA-sequencing. Fetal tissue for this experiment was obtained at day 137 of gestation then isolated and cultured as CMZ-containing far-peripheral or laminated peripheral retinospheres respectively. At day 185 equivalent gestation, the CMZ of far-peripheral retinospheres was dissected and dissociated, and laminated peripheral retinospheres were also dissociated for comparison. An equivalent number of cells from each sample were then processed with the 10x Genomics single cell RNA-sequencing platform.

When the data from the CMZ-containing far periphery were merged with that of the laminated peripheral retina we found cell clusters unique to the CMZ using Uniform Manifold Approximation and Projection (UMAP) (Fig. 3A)^19,20^. Cell types were assigned to clusters based on expression of known genes (Fig. S3A). The largest difference between cells derived from the CMZ and from the control periphery were seen in the population of early and late RPCs (red and blue) and cycling cells (purple) (Fig. 3A). These three clusters along with the neurogenic precursors (Neuro-Pre, blue-green) were subsetted and re-clustered to identify differences among progenitor cells (Fig. 3B). Cell type classes of early, late, or neurogenic progenitors were assigned based on expression of known genes in these populations (Fig. 3B, Table S1)^6^.

**Figure 3:**
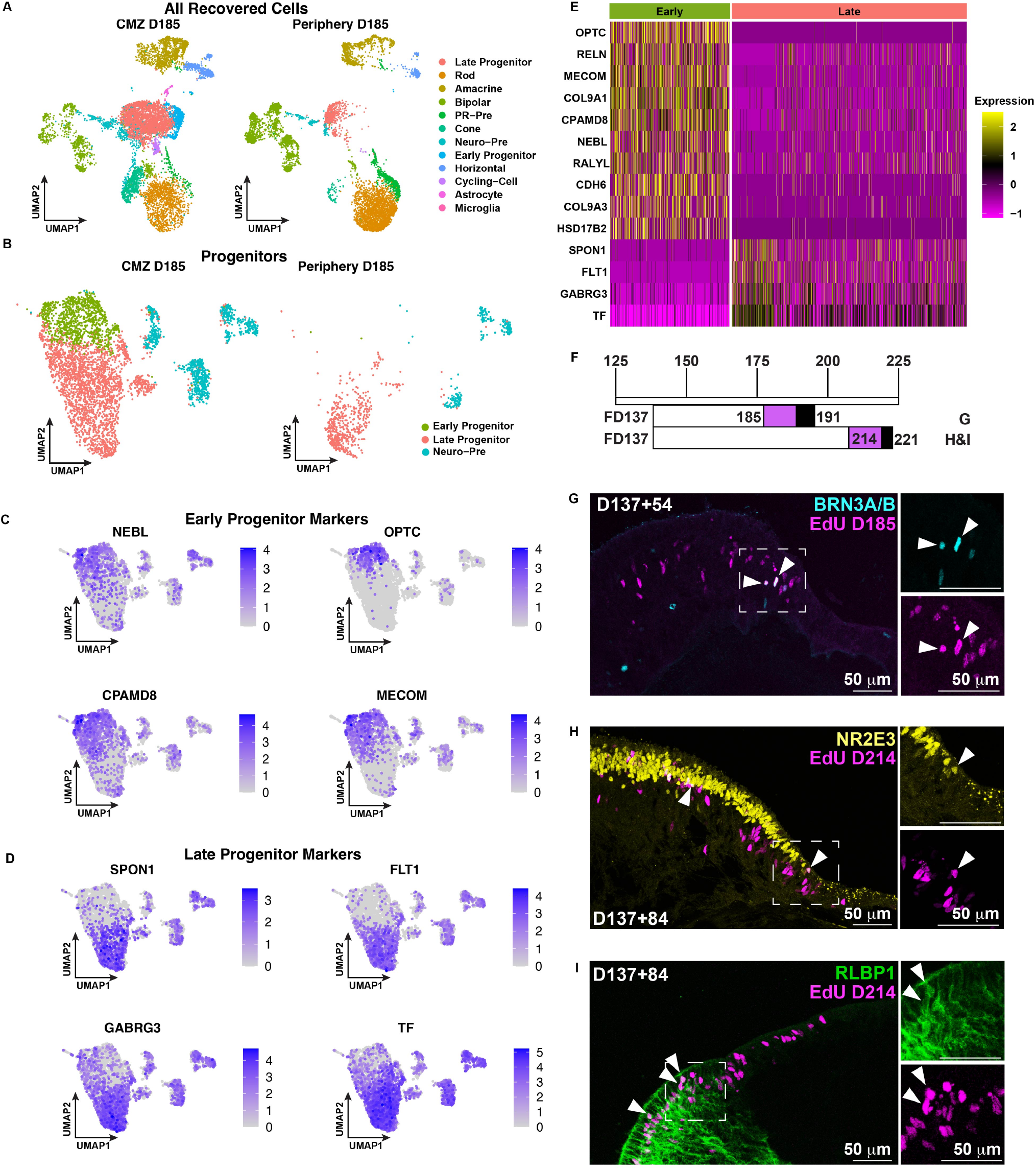
Identification of early and late retinal progenitor cells in the human fetal CMZ. **(A)** UMAP of all recovered cells from single cell RNAseq of day 185 equivalent gestation retinospheres dissected into either CMZ (left) or peripheral (right) regions. Colors indicate cell type, with corresponding ledged on the right. PR-Pre= Photoreceptor precursor, Neuro-Pre = Neurogenic precursor. **(B)** UMAP of the progenitor and cycling cells from Figure 3A, subsetted and re-clustered. CMZ (left) and peripheral (right) samples are displayed separately. **(C)** Merged UMAP of progenitor cells displayed above in B, colored by expression level of the marker genes of early progenitor cells. **(D)** Merged UMAP of progenitor cells displayed above in B, colored by expression level of the marker genes of late progenitor cells. **(E)** Heat map of select genes that are significantly differentially expressed in clusters from Figure 3B, separated by cluster. **(F)** Time course of retinosphere isolation, culture, and EdU addition. Magenta signifies EdU addition and black signifies the day of fixation. Letters “G”, “H&I” at right correspond to subsequent figures. **(G)** Retinosphere containing the CMZ isolated at 137 days gestation and cultured for 54 days, to 191 days equivalent gestational development. EdU was added on day 185 until fixing. **(H-I)** Retinosphere containing CMZ, isolated at 137 days gestation and cultured for 84 days, to 221 days equivalent gestational development. EdU was added on day 214 until fixing. **(H)** Immunostained for NR2E3 (yellow) and EdU (magenta). Arrows indicate EdU labeled NR23E positive rod, born between day 185 and day 191 equivalent gestation. **(I)** Immunostained for RLBP1 (green) and EdU (magenta). Arrows indicate EdU labeled RLBP1 positive Müller glia born between day 214 and day 221 equivalent gestation.

We then queried the top 100 differentially expressed genes between early and late progenitors of the CMZ sorted by adjusted p-value (Table S2). This list was then compared to an analogous set of genes derived from early and late progenitors from 59 and 76 day fetal retina previously profiled by single-nuclei transcriptomics in our lab^7^ (Fig. S3B). From these analyses, we found similar marker genes that have previously been described as marking early progenitors of the CMZ such as OPTC, RELN, MECOM, COL9A1, and CPAMD8^13^ (Fig. 3C, E). In addition to these previously described markers, we also found several additional genes more highly expressed in the cluster we define as early progenitors; these include NEBL, RALYL, CDH6, COL9A3, and HSD17B2 (Fig. 3C, E). Similarly, we verified that existing marker genes are present in the late progenitor cluster and further found new marker genes for late progenitors including SPON1, FLT1, GABRG3, and TF (Fig. 3D, E).

The above results indicate that the CMZ region contains cells that have a gene expression profile of both early and late progenitors. When our newly identified marker genes and other known marker genes displayed in Fig. 3E are used to create a gene module score, these modules identify early and late progenitor populations from both the D185 CMZ samples as well as the earlier D59 and D76 timepoints, indicating that the early progenitors of the CMZ are similar to those found at early time points in retinal development (Fig. S3C-D). Additionally, we found our newly identified early and late marker genes from the CMZ sample are also enriched in the D59 and D76 retinal samples’ early and late progenitors, respectively (Fig. S3E-F).

To determine if both early and late born retinal cells were generated from the proliferating population of the CMZ, we assayed for generation of ganglion cells (early), rods (late), and Müller glia (late). To identify newly generated cells, EdU was added to the culture media of CMZ containing retinospheres starting at day 185 equivalent gestation and continued for 6 days (Fig. 3F). Retinospheres were fixed at day 191 equivalent gestation, sectioned, and clickchemistry was performed for EdU, marking the newly divided cells, and immunostained for BRN3A/B, a ganglion cell specific marker (Fig. 3G). We observed colocalization of EdU and BRN3A/B, indicating that these ganglion cells were generated in the 6 days of development between day 186 and 191 equivalent gestation. In a similar experiment (Fig. 3F), we also stained for NR2E3, a marker of rod photoreceptors (Fig. 3H). We saw co-localization of NR2E3 and EdU, indicating that the later born rods are also being produced at this time. Similarly, we also observed Müller glia, the last-born cell type in the retina, labeled with both RLBP1 and EdU indicating it was produced between day 214 and 221 equivalent gestation (Fig. 3F, I). These results show that both early and late born cell types are being produced in the human CMZ at the same timepoint of approximately 190 to 220 days gestational equivalent.

### Cell cycle of CMZ progenitors is longer than that of peripheral progenitors

Previous studies have shown that cell cycle length is longer in multipotent progenitors from early stages of embryogenesis than progenitors that are committed to neuronal fate^21^. This is also observed for RPCs within the CMZ in other organisms^14,15,22^. To identify if the RPCs of the human fetal CMZ have a longer cell cycle than peripheral RPCs, we performed a timed EdU pulse experiment as described in Locker et al.^15^. EdU was added to the retinospheres for 25, 8, 6, 4, 2, or 1 hour intervals (Fig. 4A), then fixed and stained for EdU and Phospho-histone-H3 (pHH3), which labels cells in mitosis. The percentage of cells that are in mitosis and are also labeled for EdU is an estimate of the length of the cell cycle. The timepoint at which 50% of all pHH3 positive cells are also labeled with EdU indicates the halfway point of the cell cycle. We found that at the 1-hour interval, there were no pHH3+/EdU+ co-labeled cells in the CMZ containing retinospheres (Fig. 4B). However, by the 25 hour EdU addition timepoint we saw that nearly 100% of the pHH3+ cells in the CMZ were also labeled with EdU (Fig. 4C). We collated all counts of pH3+EdU+ cells from the CMZ and peripheral retinospheres and estimate that the cell cycle length of the RPCs in the CMZ is approximately 16 hours (Fig. 4D), whereas the cell cycle length of the RPCs in peripheral retina is approximately 11 hours (Fig. 4E). These data suggest that the cell cycle length of RPCs in the human CMZ is longer than that of the RPCs in peripheral retina, similar to the CMZ of other organisms^14,15,22^.

**Figure 4:**
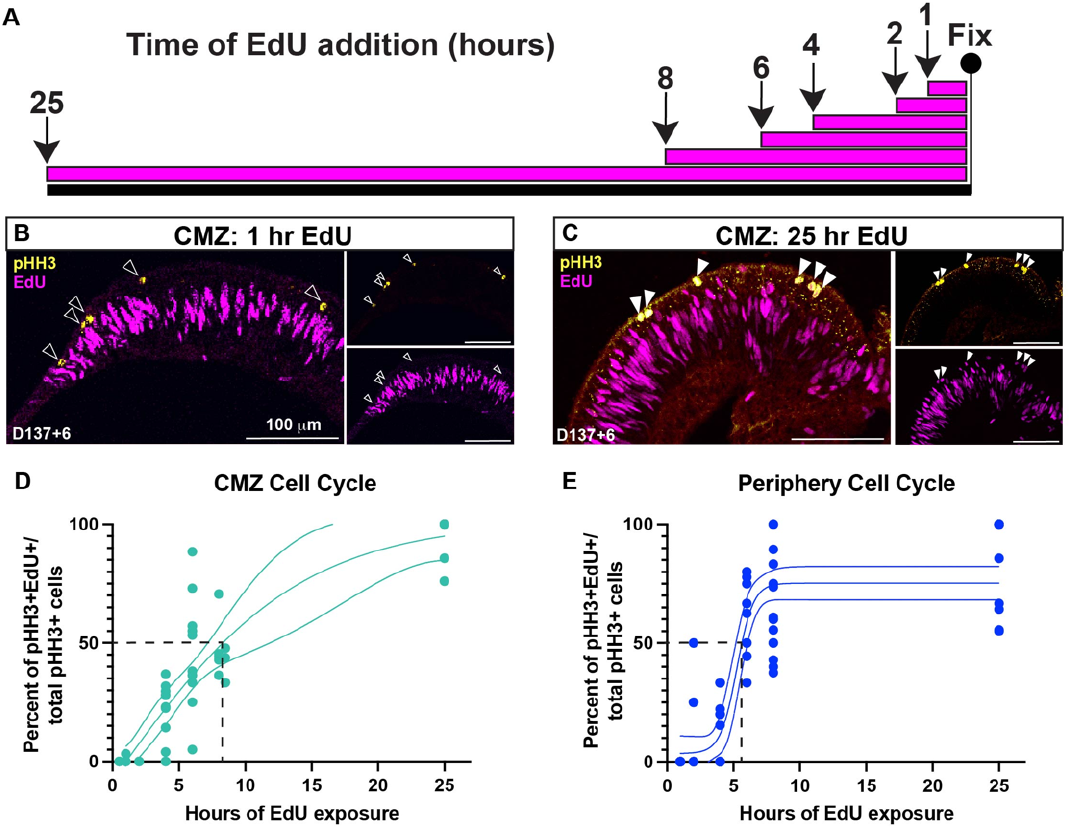
Cell cycle of CMZ progenitors is longer than that of peripheral progenitors. **(A)** Schematic of time course of EdU addition. Arrows with number indicates the hour at which the EdU was added to the culture. All retinospheres were fixed at the same timepoint after variable lengths of EdU incorporation. **(B-C)** Retinospheres containing the late proliferative zone (CMZ), isolated at 137 days gestation cultured for 6 days, to 143 days equivalent gestational development. Immunostained for phospho-histoneH3 (pHH3, yellow) and EdU (magenta). Empty arrows indicate pHH3 label without EdU. **(B)** EdU was added for one hour before fixing. 0% of EdU+ cells are co-labeled with pHH3. **(C)** EdU was added for 25 hours before fixing. 100% of EdU+ cells are co-labeled with pHH3. **(D-E)** Percent pHH3+EdU+/total pHH3+ cells over time in the CMZ, individual points are percent values per retinosphere from two separate experiments. Linear regression of data is displayed. **(D)** CMZ derived retinospheres contain 50% labeling in RPCs at 8 hours, therefore half of the cells have completed a cell cycle and the full cycle is 16 hours. **(E)** Peripheral retina derived retinospheres contain 50% labeling in RPCs at 5.5 hours, therefore half of the cells have completed a cell cycle and the full cycle is 11 hours.

### TGF-β inhibition leads to loss of late progenitor cells, however does not affect early progenitors of the CMZ

To determine how the RPCs of the CMZ respond to known regulators of cell proliferation, we treated CMZ containing retinospheres with mitogenic factors and modulators of signaling pathways. Among those we tested (Table S3), we found that SB431542, a TGF-β receptor kinase inhibitor (TGF-βi), had a differential effect on proliferation in the RPCs of the CMZ compared to those outside of the CMZ. When retinospheres containing the CMZ were treated with TGF-βi for 10 days and EdU was added for 7 days before fixing (example Fig. 5A), there was a significant decrease in the number of EdU labeled cells in the far-peripheral retina of the treatment group compared to nontreated control retinospheres (Fig. 5B-C, empty arrows); however, we saw no difference between the number of EdU positive cells in the CMZ region (Fig. 5B-C, filled arrows). Across three different treatment groups we consistently observed this pattern (Retinospheres; Control = average normalized to 1; normalized TGF-βi retinosphere = 0.2; *P* < 0.0001. CMZ; Control = average normalized to 1; normalized TGF-βi retinosphere = 8.6; *P*= ns) (Fig. 5D).

**Figure 5:**
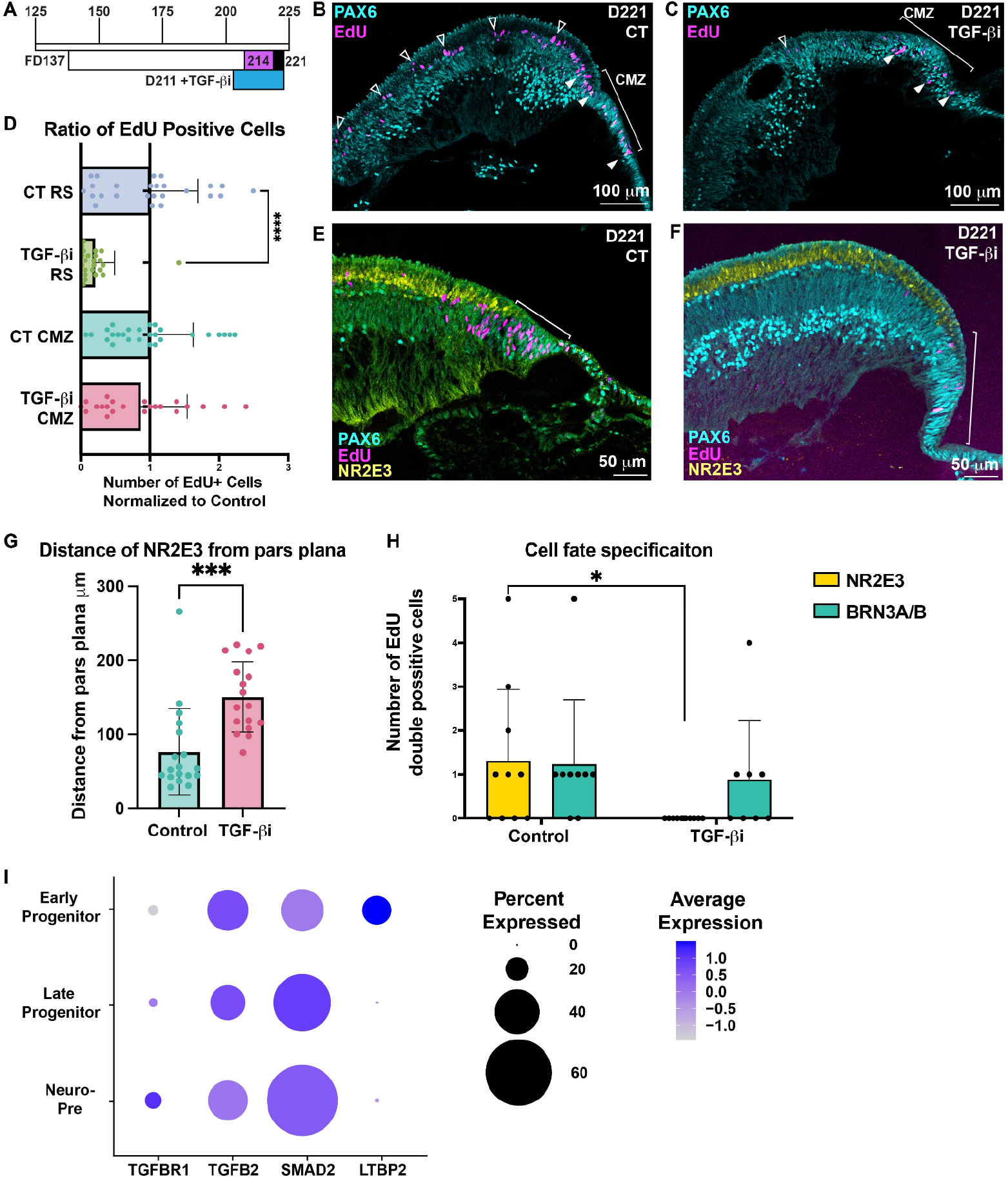
TGF-β inhibition leads to loss of late progenitor cells, however does not affect early progenitors of the CMZ. **(A)** Time course of retinosphere isolation, culture, and EdU addition. Magenta signifies EdU addition, black bar signifies final day of fixation, and blue indicates when TGF-βi was added **(B-C)** Retinospheres containing the CMZ isolated at 137 days gestation and cultured for 84 days, to 221 days equivalent gestational development. EdU was added on day 214 until fixing for a total of 7 days. Immunostained for PAX6 (cyan) and EdU (magenta). Brackets indicate CMZ region, filled arrows indicate EdU labeled cells in the CMZ, and empty arrows indicate EdU labeled cells in the far periphery of the retinosphere. **(B)** Control. **(C)** TGF-βi treated. **(D)** Ratio of the number of EdU positive cells in either the CMZ or the far periphery of the retinosphere, average number of EdU+ cells in the control is normalized to 1 for the CMZ and the far periphery of the retinosphere respectively. Mean with SD is displayed, Mann Whitney test, p < 0.0001. n= three separate experiments with a total of 27 retinospheres for control CMZ, 25 retinospheres for CT retinosphere, 22 retinospheres for TGF-βi CMZ, and 23 retinospheres for TGF-βi retinosphere. **(E-F)** Retinospheres containing the CMZ isolated at 137 days gestation and cultured for 84 days, to 221 days equivalent gestational development. EdU was added on day 214 until fixing for a total of 7 days. Immunostained for PAX6 (cyan) and EdU (magenta), and NR2E3 (yellow). Bar indicates the length measured between the pars plana and the first NR2E3 positive cell. **(E)** Control. **(F)** TGF-βi treated. **(G)** Distance from the pars plana to the first NR2E3 positive cells were measured for CMZ containing retinospheres from both control and TGF-βi treatment groups. Distance is displayed in μm. Mean with SD is displayed, unpaired t-test, p=0.0002. n= three separate experiments with a total of 18 retinospheres for control and 17 retinospheres for TGF-βi treatment. **(H)** Quantification of cell fate specification in retinospheres. Double positive EdU and NR2E3 cells are quantified for control (n=3 experiments, 10 retinospheres and 15 total sections) and TGF-βi treatment (n=3 experiments, 10 retinospheres and 17 sections); Mann Whitney T-test, p=0.0108. Double positive EdU and BRN3A/B cells are quantified for control (n= 2 experiments, 10 retinospheres and 14 sections) and TGF-βi treatment (n=2 experiments, 8 retinospheres, and 14 sections); Mann Whitney T-test p=ns. Mean with SD is displayed. **(I)** Average expression level and percent of cells expressing genes involved in the TGF-β signaling pathway for clusters displayed in Fig. 3B.

In addition to the effects on EdU incorporation, there also appeared to be changes in the rate of rod differentiation: the distance between the pars plana and the first NR2E3 positive cell was greater in TGF-βi treated retinospheres than in controls (Control = 84.5 μm; TGF-βi = 156.9 μm; *P* = 0.0002) (Fig. 5 E-G). To further evaluate the rate of rod differentiation we labeled the dividing cells in the spheres for 7 days with EdU and then co-labeled with markers of ganglion cells (BRN3A/B) and rods (NR2E3). We then counted the number of cells that were double positive for EdU and one of these markers, indicating that they were newly differentiated in the past 7 days of EdU treatment. In control CMZ-containing retinospheres, we saw on average 1 newly differentiated ganglion cell, and an average of 1 newly differentiated rod per retinosphere (Fig. 5H). However in TGF-βi treated CMZ-containing retinospheres we see on average one newly differentiated ganglion cell per retinosphere and no newly differentiated rods (Control = 1.6, TGF-βi = 0; *P* = 0.01) (Fig. 5H). These results suggest that TGF-βi affects the rate of rod differentiation, and selectively causes progenitors in the far peripheral retina to exit the cell cycle.

The observation of differential responses to TGF-βi signaling prompted us to interrogate the expression levels of proteins involved in the TGF-β signaling pathway. When comparing the gene expression levels of early progenitors, late progenitors, and neuro-precursors in the single-cell RNA-seq dataset presented above (Fig. 3B), we find that early progenitors have lower levels of the receptor TGFBR1, lower levels of SMAD2, a co-factor for TGF-β signaling, and significantly higher amounts of LTBP2 (*P* < 0.0001), which inhibits TGF-β signaling (Fig. 5I). Interestingly, early progenitors also have slightly elevated levels of TGFB2, the most highly expressed ligand of this family found in the retina. Taken together, these data suggest that early progenitors may be less responsive to TGF-β signaling than late progenitors or neurogenic precursors, potentially accounting for their lack of response to TGF-βi treatment.

## Discussion

The CMZ of frogs and fish continues to grow throughout their adult lives and contributes to regenerating retinal cells upon injury^14,23,24^. Additionally, some growth post-hatch has also been observed in the chick retina^8,25,26^. There has been debate as to whether humans and other mammals maintain this proliferative CMZ region^27-31^; however, it is generally accepted that the human retina does not have a defined stem cell zone within the CMZ that contributes to the laminated retina in adulthood^9,10,32,33^. Recent single cell RNA-seq analysis has observed a CMZ-like region is present in the fetal human retina at 91 days gestation, however a gap remains in our knowledge regarding how long the human CMZ maintains proliferation during development. Here, we show that human fetal retina has a CMZ-like region, similar to frogs and fish. Additionally, we show that this region persists in late stages of human development, and maintains a zone of RPCs that is capable of giving rise to all the cell types within the retina.

To date, studies of the fetal human CMZ have been limited to primary fixed tissues, mostly from early gestational timepoints and a limited number of samples over day 150 of gestation^13^. In this study, we add to these prior observations with investigation of human gestational time points up to 238 days, as well as rhesus monkey tissues isolated at day 145 of gestation, the equivalent of 260 days humans gestation. These observations add to our understanding that the human CMZ continues to proliferate up until at least day 238 of gestation in humans, and close to birth in rhesus monkeys (Fig. 6A).

**Figure 6:**
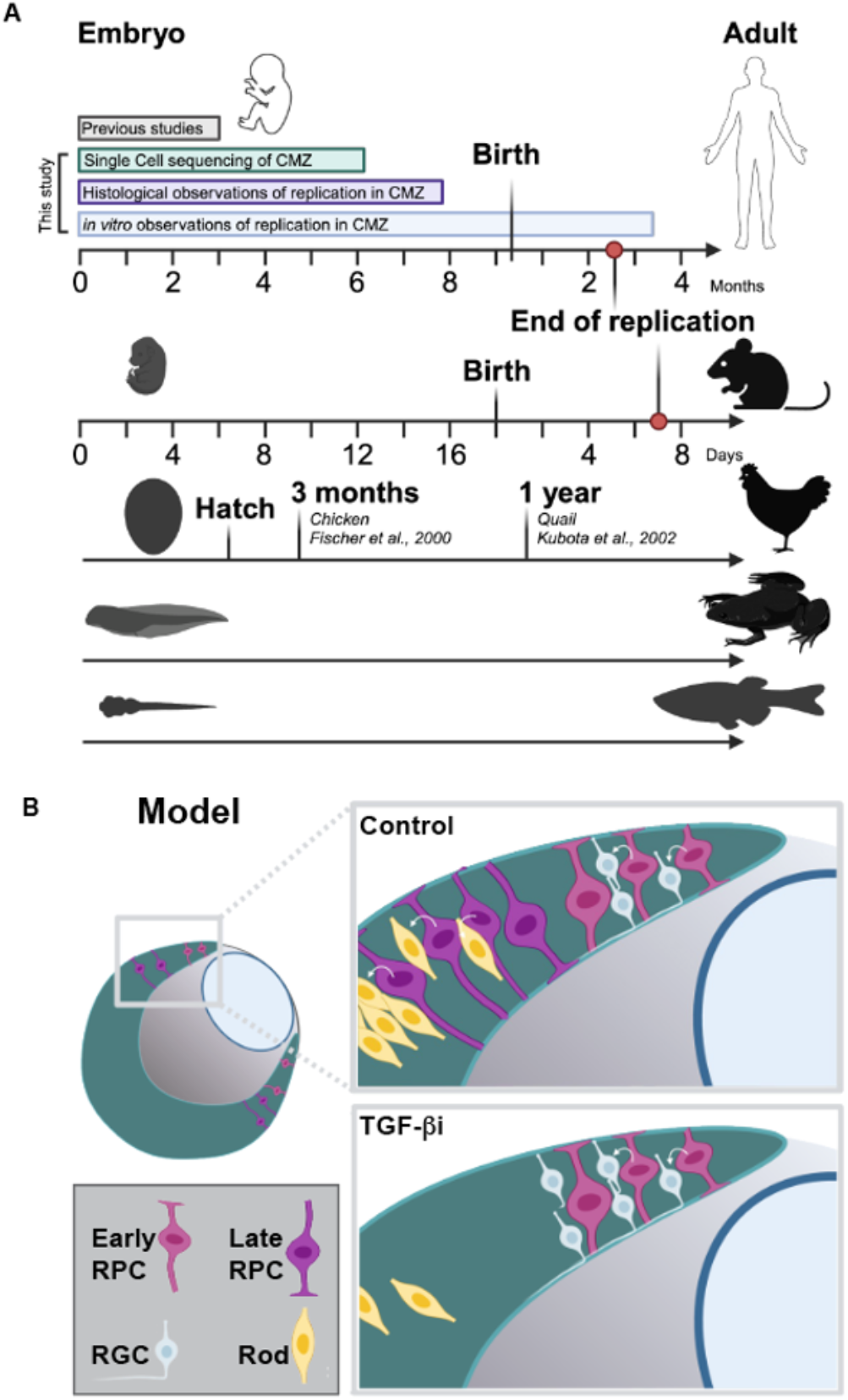
Timeline of observations of developing human CMZ greatly extended. **(A)** Diagram of timing of CMZ development across organisms. Human development displayed in months. Previous studies performing spatial-RNA-seq in the CMZ have reached 91 days gestation^13^. In this study we provide scRNA-seq at the late timepoint of 185 days equivalent gestation, histological observations of 238 day gestation tissues, and *in vitro* observations of replication in the CMZ until 2.5 months postnatal development. Mouse development displayed in days. Chick and other avian CMZ proliferation has been observed at 3 months^8^ and 1 year^55^, respectively. Frog and zebrafish continue to proliferate until adulthood. **(B)** Model for how TGF-βi is affecting the progenitor cells of the CMZ verses those of the retinosphere differently. Early progenitors, located at the tip of the CMZ, give rise to early cell types such as ganglion cells. Late progenitors, located further out in the far-periphery of the retinosphere, give rise to late born cell types such as rods. Upon addition of TGF-βi late progenitors exit the cell cycle and no longer differentiate rods, while early progenitors are not affected by the inhibitor. In this way, cells are continually added through division by early progenitors, but not by late progenitors, thereby increasing the distance between the pars plana and the first NR2E3 positive rod.

Previous studies of the human retina’s capacity for proliferation have mainly focused on dissociated retinal cultures derived from readily accessible early gestational time points. Due to the difficulty of capturing the small CMZ region it is unlikely that it was included in the cultures^34,35^. In our study, we expanded our previously described retinosphere culturing system^5^ to isolate and culture the CMZ of human fetal tissue. This method allows for further development of the retina while maintaining specific spatial identities^5^ (Fig 1. D-I). With this system we were able for the first time to observe the amount and localization of proliferation at late gestational time points in the human CMZ, and provided evidence that this proliferation may continue at postnatal timepoints (Fig. 2H, 6A). Further studies in primary tissues directly from postnatal time points are needed to confirm the presence of these proliferating cells in the CMZ postnatally.

Other studies have conducted both bulk and single cell transcriptomic profiling of retinal tissue from similar timepoints to look at global gene changes in retinal development; however, due to the difficulty of dissecting this region and the small size of the CMZ compared to the rest of the retina, it is not surprising that these studies have not observed early RPCs past day 105 of gestation^6,13,36^. For accurate assessment of the RPCs present in the human CMZ, this region must be specifically isolated and enriched for, or observed through spatial transcriptomics. Prior to this study, the furthest developmental timepoint in which this region has been directly observed was via spatial RNA-seq up to day 91 of gestation^13^. Here we provide single cell transcriptomic profiles of mechanically isolated CMZ vs. peripheral retina from day 185 of development, past the time points that are normally accessible in human gestation, and show for the first time that at the late gestational time point of D185 the human CMZ maintains early RPCs (Fig. 6A).

In keeping with the observation that the human CMZ maintains both early and late RPCs, we also showed for the first time that this region continues to generate both early born (ganglion) and late born (rod and Müller glia) cell types until at least day 220 of equivalent gestational development (Fig. 3F-I). These findings are consistent with previous studies in the mouse, where ganglion cells are observed to be generated at postnatal timepoints, equivalent to late gestational time points in human development^12^. This is particularly interesting as one model for the temporal progression of RPCs is based on the assumption that RPCs progress through different windows of competency over time^2,4,37^. The fact that ganglion cells and Müller glia are being generated at the same time in such close proximity to one another suggests that there may be spatial cues, perhaps from the RPE or the CMZ, that RPCs are responsive to in order to govern their early and late RPC identity. This also implies that temporal identity factors can be expressed in a more rapid period of time in the CMZ than the longer time scales observed during central and peripheral retinal development.

The cell cycle length is longer in multipotent progenitors from early stages of embryogenesis, or more “stemlike” cells, than progenitors that are committed to a neuronal fate^21^. Here, we show that, similar to other model organisms^14,15,22^, the RPCs of the CMZ region have a longer cell cycle than those of the laminated peripheral retina (Fig 4D-E). As these experiments were completed at later time points in development, most of the proliferating cells in this region are late RPCs. It is possible that we are seeing a difference in the cell cycle length between early and late RPCs, rather than specifically a unique characteristic of RPCs within the CMZ. Further studies would be needed to define the cell cycle kinetics of early and late peripheral RPCs in addition to those of the CMZ to delineate between these two possibilities.

To further test the hypothesis that the signaling environment can influence RPC identity and differentiation, we treated with several small molecules and screened for effects (Table S3). In other model organisms that contain a proliferative CMZ after birth, a number of signaling pathways have been identified to govern the proliferation and maintenance of these retinal stem cells, including Yap, Wnt2b, Shh, insulin, EGF, FGF, NF2, and BMP^11,14,22,38-45^. The only small molecule we identified from those we tested that had a measurable effect on proliferation was TGF-β inhibition by treating with SB431542.

We observe that TGF-β inhibition preferentially forced late RPCs to exit the cell cycle, as seen by a significant decrease in EdU positive cells in the far-peripheral regions of the retinospheres (Fig. 5D), whereas early progenitor cells were maintained in the CMZ. This is particularly surprising given the wealth of literature indicating that TGF-β signaling leads to neuronal differentiation, therefore suggesting that TGF-β inhibition would increase proliferation^46^. Indeed, this has been observed by our group in dissociated rat retinal cultures where proliferation was restored in late progenitors treated with TGF-β inhibitor TGF-βRII-Fc fusion protein^47^. However, our group has also observed the opposite phenomenon of TGF-β3 treatment leading to more proliferation in rat retinal cultures^48^. It is possible that different retinal regions respond differently to TGF-β signaling, similar to astrocytes of the brainstem versus those of the forebrain, which respond differentially to TGF-β3^49^. It would be interesting to test the effect of TGF-β inhibition on early and late progenitors in more central retinal regions to understand if the same principals we observed in the CMZ hold true in other retinal regions.

Taken together, we propose the following model: in control tissues early RPCs are located at the distal tip of the CMZ where they produce ganglion cells as well as other RPCs. Moving away from the CMZ, late RPCs are present that are creating rods and other late born cell types such as Müller glia (Fig. 3G, H). Upon treatment with TGF-βi, late RPCs exit the cell cycle and are no longer present to generate rods (Fig. 6B). However the early RPCs continue to proliferate, generating differentiated cells and increasing the distance between the pars plana and the first NR2E3 expressing cell (Fig. 6B). We propose that this is caused by differential capacities to respond to TGF-β signaling, as early progenitors have lower levels of the receptor TGFBR1, lower levels of SMAD2, a co-factor for TGF-β signaling, and higher amounts of LTBP2, which inhibits TGF-β signaling (Fig. 5I). In conclusion, we provide here a new method for isolating and culturing fetal CMZ of the developing human retina. This system has allowed for the first functional studies of this region at late developmental time points, and identification of the timeline of proliferation and senescence of RPCs in the human retina. We further define the cell cycle length of the RPCs of the CMZ, and show that it is longer than those of the peripheral retina. Additionally, this method has provided insight into the requirement for proper TGF-β signaling to maintain late RPCs, however it is dispensable for maintenance of early RPCs. Taken together, these studies highlight the utility of this region for further study of RPC maintenance, as well as temporal and spatial signaling factors that influence cell fate specification in the human retina. A better understanding of the kinds of RPCs that are present in the human retina and what governs their proliferation will aid in our understanding of basic retinal development, as well as give insights into regenerative therapies to restore vision in blinding diseases such as glaucoma and macular degeneration^50-52^.

## Acknowledgements

We would like to thank all the members of the Reh lab, the Bermingham-McDonogh lab, and Rachel Wong for their valuable comments on the manuscript. This research was supported by Foundation Fighting Blindness (TA-RM-0620-0788-463 UWA to T.A.R.), the Damon Runyon Cancer Research Foundation (DRG-#32-20 to K.C.E.), and a Hanna H. Gray Fellows Program Award from the Howard Hughes Medical Institute (Grant #GT15994 to K.C.E.), and the Generalitat Valenciana (APOSTD/2020/245 to I.O.L.). Illustrations were created with BioRender.com.

## Author contributions

Conceptualization, T.A.R., K.C.E, and I.O.L; Methodology, K.C.E, and I.O.L; Software, K.C.E.; Formal Analysis, K.C.E.; Investigation, K.C.E, I.O.L, S.J.E., J.W., S.M.S., S.P., G.W.D., D.H.,; Resources, I.G., A.L.T., T.A.R.; Writing – Original Draft, K.C.E.; Writing –Review & Editing, K.C.E. and T.A.R.; Funding Acquisition, T.A.R., K.C.E., I.O.L.; Supervision, T.A.R., and K.C.E.

## Supplemental information

**Document S1**. Figures S1–S3 and Table S3 and Table S4.

**Table S1:** Table_S1_RPC_Genes.xlsx, excel file containing additional data too large to fit in a PDF, related to Figure 3.

**Table S2**. Table_S2_RPC_MarkerGenes.xls, excel file containing additional data too large to fit in a PDF, related to Figure 3.

**Table S3**

**Table S4**

## Materials and Methods

### Fetal retina tissue

Human fetal retinal tissue was obtained from the Birth Defects Research Laboratory at University of Washington using an approved protocol (UW5R24HD000836). Tissues had no identifiers, and ultrasounds along with physical characteristics such as fetal foot length and crown-rump were used to estimate the gestational age^53^.

### Rhesus monkey tissue

All animal procedures conformed to the requirements of the Animal Welfare Act and protocols were approved prior to implementation by the Institutional Animal Care and Use Committee (IACUC) at the University of California at Davis. Healthy adult female rhesus monkeys (Macaca mulatta) were time-mated and identified as pregnant using established methods54. Normal fetal growth and development stage were confirmed by ultrasound during gestation54. Pregnancy in the rhesus monkey is divided into trimesters by 55-day increments: 0-55 days gestational age represents the first trimester, 56-110 days represents the second trimester, and 111-165 days the third trimester (term 165 ± 10 days)^18^. Dams were scheduled for hysterotomy for fetal tissue collection and were returned to the breeding colony post-hysterotomy.

The fetal eyes were collected into cold media (DMEM supplemented with 10% FBS) then extraneous tissue was removed and the retina was incubated in oxygenated media for 90 min at room temperature. Samples were then fixed in modified Carnoy’s fixative (ethanol, formaldehyde, and acetic acid) overnight at 4°C, dehydrated in stepwise ethanol/water solutions, cleared with xylene, and embedded in paraffin blocks.

### Retinospheres (*in vitro* fetal retinal cultures)

Retinospheres were made and cultured as described in Sridhar et al., 20205. In brief, the fetal retina was dissected into 300–400-micron pieces with dissecting scissors. These were then maintained in RDM (Retinal differentiation medium, 46% DMEM, 46% DMEM/F12, 2% B27, 1% Pen Strep and 5% FBS) in low-adhesion 6 well plates or 10cm dishes. To create region-specific retinospheres such as CMZ containing retinospheres, target regions of the fetal retina were mechanically dissected, then cut into retinospheres. Retinospheres were cultured using approved BSL2 methods and cells were maintained at 37C in 5% CO2 conditions. The following small molecules were added to the cultures: EdU (10 uM, Click-iT EdU Assay, Invitrogen), SB431542 (5uM, Selleckchem).

### Tissue dissociation for 10x

The CMZ was dissected from the retinospheres or peripheral retinospheres were isolated and dissociated using Accutase and DNAse for 20-30 minutes on a nutator at 37C, with gentile pipetting to break up cells every 7 minutes.

Cell pellets were resuspended in PBS containing 0.04% bovine serum albumin and filtered through a 35µm cell strainer (Fisher, 08-771-23) to remove cell clumps. Cells were then counted with a hemocytometer. 8,000-10,000 cells of the sample were then used as input into the 10X protocol. GEM generation, reverse transcription, cDNA amplification, and library construction steps were performed according to the standard 10x workflow for Single Cell 3 v3.1 Dual Index Gene Expression kit (Dual Index Kit TT Set A 96 rxns, 10x Genomics, 1000215).

### scRNaseq analysis and data processing

Seurat analysis was performed in R using Seurat v5.0.3, UMAP (Becht et al., 2018)^20^, ggplot2, and dplyr. The raw counts were normalized, scaled, and variable features were identified. Datasets combined in Seurat were integrated by performing CCA on the combined datasets, and reduced into a lower dimensional space using dimensional reduction (Butler et al., 2018)^19^. Clustering resolution for all datasets were set so clusters matched well with known cell type markers (Fig. S2A). Cell types were assigned to clusters in the merged objects.

### Sectioning and immunofluorescence

Fetal retinas at younger ages (D50-Day70) were fixed without lens removal with 4% paraformaldehyde (1 hr). Older retinas were first fixed with a small opening in the lens for 40 minutes at 4C, followed by lens removal and overnight fixation in 4% paraformaldehyde at 4C. Retinospheres were fixed in 4% paraformaldehyde, 1X PBS, and 5% sucrose at 4C for 45 minutes followed by 3 washes in 1X PBS, 5% sucrose. All samples were then transitioned into 10%, 20% and 30% solutions of sucrose, followed by an additional 30% sucrose +OCT step for the human retina. These were then embedded in OCT and cryosectioned at 14-16 mm.

Immunostaining was performed as previously described in Hoshino et al., 201736. Briefly slides were blocked in 0.5%Triton/2% horse serum for an hour, followed by primary antibody incubation in 0.5%Triton/2% horse serum at 4C overnight. The next day, the slides were washed 3 times in 1X PBS, followed by secondary antibody incubation in 0.5%Triton/2% horse serum for an hour at room temperature. For EdU labeling, cells were first fixed for 5 min in 4% PFA after secondaries were added, then washed 3x in PBS for 5 min each. Slides were then incubated for 30 min at RT with the Click-it solution (Click-iT EdU Assay, Invitrogen) and then washed three times with PBS before being mounted. Slides were then mounted using Fluromount.

NHP tissues were embedded in paraffin as detailed above, and sectioned at 8-16 microns. Slides were then deparaffinized and rehydrated my immersing the slides through the following solutions for 5 min each: 1) Xylene, three washes. 2) 100% Ethanol, two washes. 3) 95% Ethanol, one wash. 4) 70% Ethanol, one wash. 5) 50% Ethanol, one wash. 6) DI water, one wash. Slides were then marked with a hydrophobic barrier pen, and permeabilized in PBS + 0.3% triton for 15 min. Heated antigen retrieval was preformed by microwaving samples in 0.1 M Citric Acid, pH 6 for 30 seconds until boiling, rest 5 min, and then heated to boil once again. Slides were then washed in PBS for 5 min. Blocking and antibody additions lited in Table S4 were performed as previously described for cryosectioned tissues.

### Microscopy

Brightfield images were obtained at 4x magnification using the Zeiss Axio Observer D1. Confocal images were obtained using the Zeiss LSM 990. Image processing was done in ImageJ and Adobe Photoshop.

### Data and code availability

The datasets generated within this study will be available on GEO upon publication in per reviewed journal.

## Supplementary Information

**Table S1: Table_S1_RPC_Genes.xlsx**, excel file containing additional data too large to fit in a **PDF**, related to Figure 3^*^

**Table S2: Table_S2_RPC_MarkerGenes.xls**, excel file containing additional data too large to fit in a **PDF**, related to Figure 3^*^

* Available upon publication in peer-reviewed journal, or by corespondance with T.A.R. –

**Table S3:**

| Molecule | Activity | Pathway | Effect |
| --- | --- | --- | --- |
| **SB431542** | TGFB inhibitor | TGFB signaling | Fewer late RPCs |
| **MGH-CP1** | TEAD inhibitor | TEAD signaling | none |
| **CHIR 99021** | Wnt Activator | Wnt Signaling | none |
| **BMP4** | signaling molecule | BMP4 signaling | none |
| **EGF** | signaling molecule | EGF signaling | none |
| **DBZ** | inhibits gamma-secretase | Notch pathway | none |
| **Cyclopamine** | Hedgehog signaling inhibitory | SHH pathway | Peripheral progenitors are decreased |
| **SAG** | Smoothened agonist | SHH pathway | none |

**Table S4:**

| Antibody | Source | Catalog # | Dilution |
| --- | --- | --- | --- |
| BRN3A/B | SantaCruz | SC-6026 | 1:300 |
| PAX6 | BioLegend | 901001 | 1:300 |
| VSX2 | Exalpha Biologicals | X1180P | 1:300 |
| VSX2 | SantaCruz | Sc-365519 | 1:300 |
| OTX2 | ABCAM | 21990 | 1:300 |
| CCND1 | ABCAM | AB134175 | 1:100 |
| RLBP1 | ABCAM | ab15051 | 1:200 |
| DAPI | Invitrogen | D1306 | 3 mM final |
| HUC/D | Life Technologies | A-21271 | 1:200 |
| KI67 | BD Pharmigen | 550609 | 1:250 |
| SOX2 | Merck | AB5603 | 1:250 |
| NR2E3 | Perseus Proteomics | PP-H7223-00 | 1:100 |
| pHH3 | NOVUS Biologicals | NB600-1168 | 1:300 |
| Donkey anti-goat 568 | Invitrogen | A11057 | 1:300 |
| Donkey anti-goat 488 | Jackson | 705-545-147 | 1:300 |
| Donkey anti-mouse 488 | Jackson | 715-605-150 | 1:300 |
| Donkey anti-mouse 568 | Invitrogen | A10037 | 1:300 |
| Donkey anti-mouse 647 | Invitrogen | A32787 | 1:300 |
| Donkey anti-rabbit 488 | Invitrogen | A21206 | 1:300 |
| Donkey anti-rabbit 568 | Invitrogen | A10042 | 1:300 |
| Donkey anti-rabbit 647 | Invitrogen | A31573 | 1:300 |
| Donkey anti-sheep 488 | Invitrogen | A11015 | 1:300 |

**Fig. S1:**
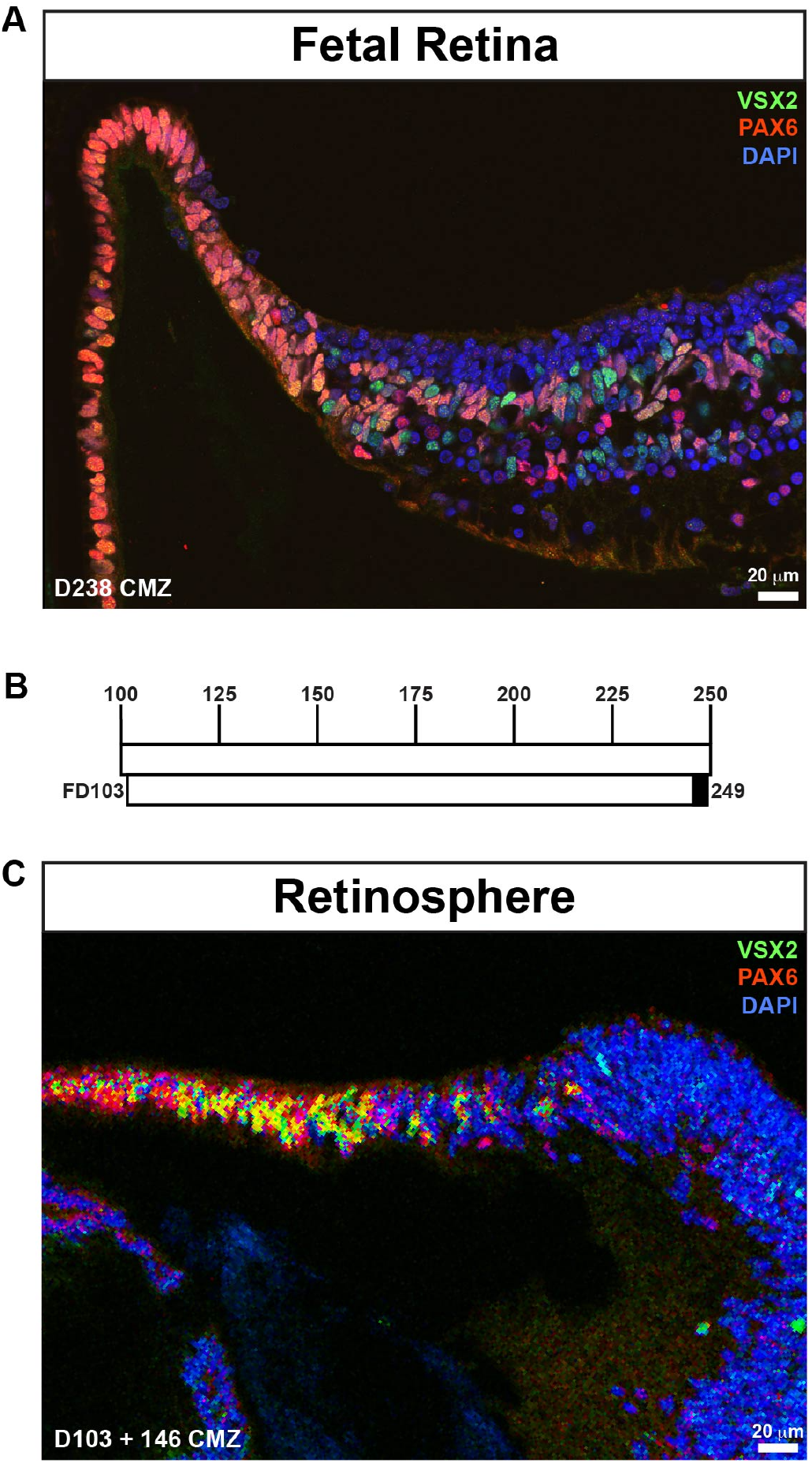
Related to Figure 1. **(A)** Fetal retina, isolated and fixed at day 238 of gestational development. Scale bars are 20 μm. Immunostaining in the CMZ for VSX2 (green), PAX6 (red), and DAPI (blue). **(B)** Time course of retinosphere isolation, culture, and fixation for retinosphere displayed in C. Black bar signifies final day of fixation. **(C)** Retinosphere containing the late proliferative zone (CMZ), isolated at 103 days gestation and cultured for 146 days, to 249 days equivalent gestational development. Immunostaining in the CMZ for VSX2 (green), PAX6 (red), and DAPI (blue).

**Figure S2:**
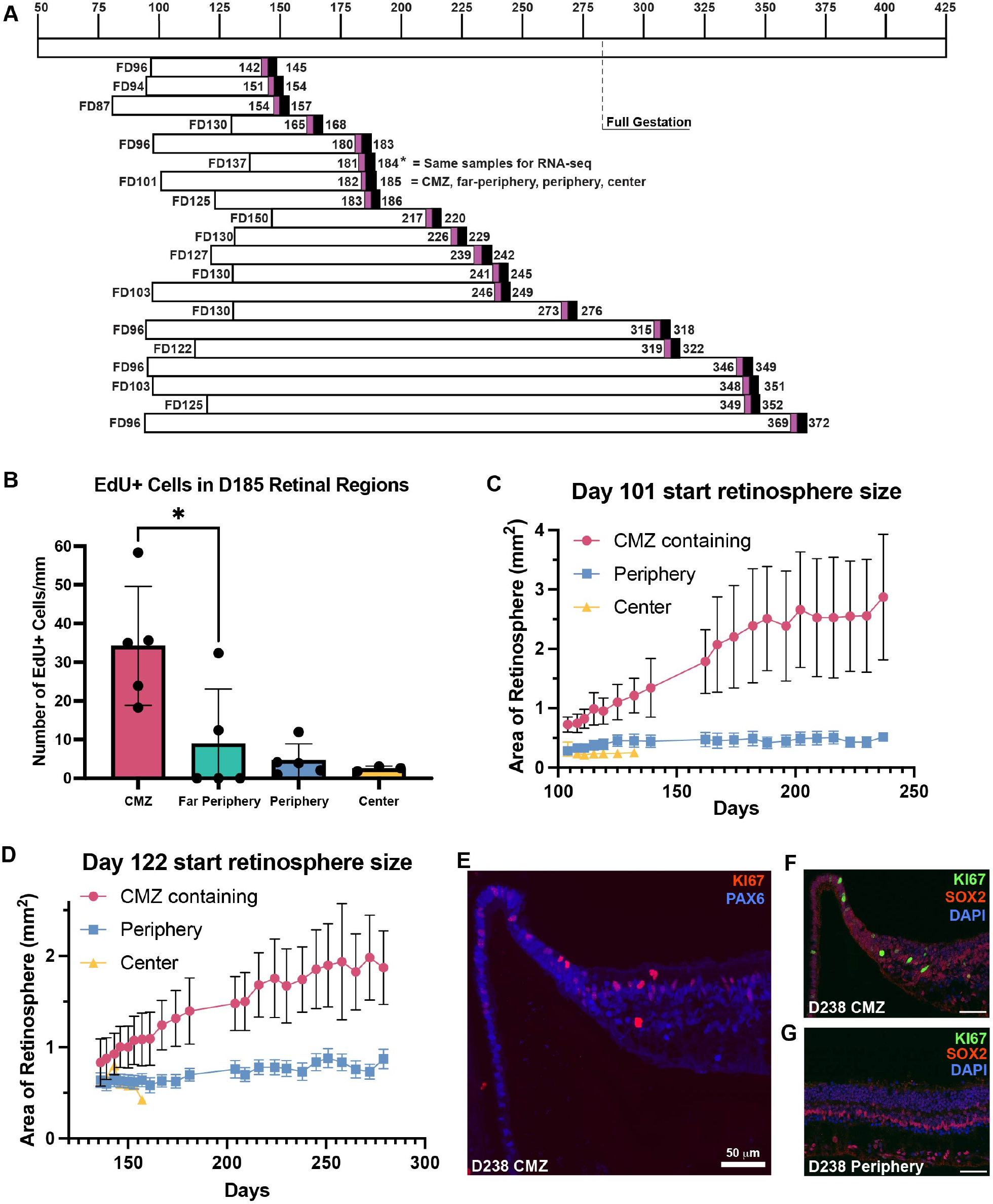
Related to Figure 2. **(A)** Time course of retinosphere isolation, culture, and EdU addition for experiments quantified in Fig. 2H. Magenta signifies EdU addition, black bar signifies final day of fixation. **(B)** Retinospheres were isolated at day 101 of fetal development from center, periphery, or the CMZ. EdU was added to the cultures for the last three days of development, from day 182 until day 185 when the retinospheres were fixed and stained for EdU. Bars show the average and SD of number of EdU+ cells per mm. Unpaired T-test of CMZ vs. FP, p = 0.0269. **(C-D)** Area of retinospheres over time. Retinospheres were isolated from the CMZ, periphery, or central retina and brightfield imagines were obtained while being cultured. Two-dimensional area of the retinosphere was then measured using ImageJ. Center and peripheral derived retinospheres largely stay the same size in culture, whereas the CMZ derived retinospheres continue to increase in size until day 275 of equivalent gestation. **(C)** Retinospheres were isolated at day 101 of fetal development. SEM is displayed, Kruskal-Wallis Multiple Comparison tests; CMZ vs. Periphery, p <0.0001, CMZ vs. Center, p < 0.0001, Periphery vs. Center, p = ns. **(D)** Retinospheres were isolated at day 122 of fetal development. SEM is displayed, Kruskal-Wallis Multiple Comparison tests; CMZ vs. Periphery, p <0.0001, CMZ vs. Center, p < 0.0001, Periphery vs. Center, = ns. **(E-G)** Fetal retina, isolated and fixed at day 238 of gestational development. Scale bars are 50 μm. **(E)** Immunostaining in the CMZ for KI67 (red) and PAX6 (blue). **(F)** Immunostaining in the CMZ for KI67 (green), SOX2 (red), and DAPI (blue). **(G)** Immunostaining in the peripheral retina for KI67 (green), SOX2 (red), and DAPI (blue).

**Figure S3:**
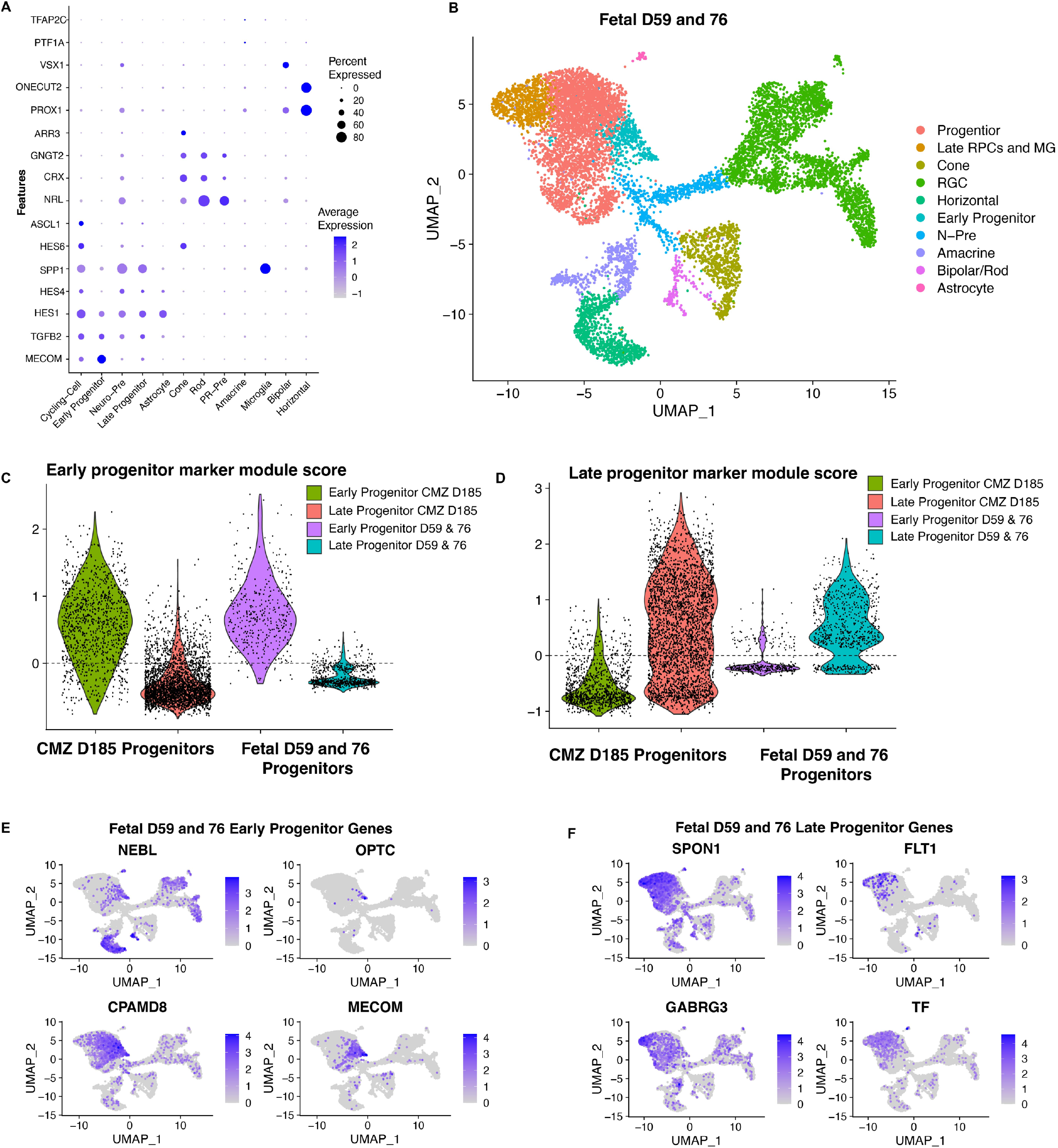
Related to Figure 3. **(A)** Average expression level and percent of cells expressing marker genes used to define clusters for UMAP of all recovered cells from single cell RNA-seq of day 185 equivalent gestation retinospheres dissected from CMZ and peripheral regions displayed in UMAP in figure 3A. **(B)** UMAP of all recovered cells from previous single cell RNA-seq of fetal days 59 and 76^7^. Colors indicate cell type, with corresponding ledged on the right. RPC = Retinal Progenitor cell, MG = Müller glia, N-Pre = neurogenic precursor. **(C)** Early progenitor marker score module consisting of early genes from Fig. 3E, displayed for both early and late progenitor clusters from the CMZ day 185 progenitors or the fetal day 59 and 76 progenitors. The early cluster from both datasets are enriched for early markers. **(D)** Late progenitor marker score module consisting of early genes from Fig. 3E, displayed for both early and late progenitor clusters from the CMZ day 185 progenitors or the fetal day 59 and 76 progenitors. The late cluster from both datasets are enriched for late markers. **(E)** UMAP of all cells from fetal day 59 and 76 displayed above in B, colored by expression level of the genes NEBL, OPTC, CPAMD8, and MECOM, which are marker genes of early progenitor cells. **(F)** UMAP of all cells from fetal day 59 and 76 displayed above in B, colored by expression level of the genes SPON1, FLT1, GABRG3, and TF, which are marker genes of late progenitor cells.

## Notes

### Competing Interest Statement

The authors have declared no competing interest.

